# Cell-type-resolved RNP topologies reveal dynamic structural mechanisms of splicing and therapeutic targets

**DOI:** 10.64898/2026.05.17.725762

**Authors:** Jianhui Bai, Kongpan Li, Wei Zhang, Minjie Zhang, Grant Shoffner, Willem A. Velema, Jianfu Chen, Feng Guo, Zhipeng Lu

## Abstract

Resolving RNA conformations in native ribonucleoprotein (RNP) complexes remains a fundamental challenge. Here, we introduce spatial hydroxyl acylation reversible crosslinking with immunoprecipitation (SHARCLIP) to simultaneously capture RNA-RNA, RNA-protein and protein-protein contacts in cells. SHARCLIP profiling of HNRNPC-associated RNA conformations established a global phased map of ribonucleosomes, resolving a decades-old debate on heterogenous nuclear (hn)RNP assembly. We built a dynamic structural atlas across seven cell lineages for >10,000 RNAs, generating ~200 million contacts, and identifying millions of dynamic loops, steric blockers and conformational switches that control splicing outcome. Deciphering the structural logic of mutually exclusive exons (MXEs) enabled rational design of structure-breaking and stabilizing antisense oligonucleotides (ASOs). We demonstrate effective isoform swapping in 12 genes linked to genetic disorders. SHARCLIP provides a comprehensive roadmap for cellular RNA structural biology and structure-guided RNA therapeutics.

## Main Text

Alternative splicing (AS) generates proteomic and organismal complexity through a regulatory code traditionally defined by linear cis-elements and trans-acting RNA-binding proteins (RBPs) (*1-3*). However, RNA naturally folds into intricate structures within ribonucleoprotein (RNP) complexes, a critical layer of regulation that remains largely unmapped. This structural blind spot has profound clinical consequences. Up to one-third of pathogenic mutations impact splicing, driving common neurodegeneration, muscular dystrophies, and cancer, and potentially thousands of rare genetic disorders (*4, 5*). While targeting RNA with antisense oligonucleotides (ASOs) and small molecules has emerged as a powerful therapeutic modality, current ASO drug development is hindered by a narrow focus on the linear sequence and reliance on blind screening, leaving the vast structural landscape beyond reach (*6-8*).

Dissecting the multivalent RNP complexes in cells is a formidable challenge (**Fig. 1A**). CLIP-like methods map RNA-protein contacts without structural context *(9-11)*. Chemical probing profiles one-dimensional RNA reactivity for local conformations (*12, 13*). Recent proximity-ligation methods rely on crosslinkers such as UV, psoralens and formaldehyde that suffer from strong bias, low efficiency, and irreversible damage (*12, 14*). To address this challenge, we developed SHARCLIP by integrating spatial hydroxyl acylation reversible crosslinking (SHARC) (*15*) with immunoprecipitation to simultaneously capture multivalent RNP interactions. Applying SHARCLIP, we resolved the architecture of cellular hnRNP particle (*16*), establishing phased ribonucleosomes as the unit of nuclear RNA packaging, analogous to nucleosomes in chromatin (*17*). By constructing a dynamic structural atlas across seven cell lineages, we uncovered the topological rules governing splicing and demonstrated proof of concept for structure-guided RNA therapeutics for a large number of splicing-related genetic disorders.

**Fig. 1.**
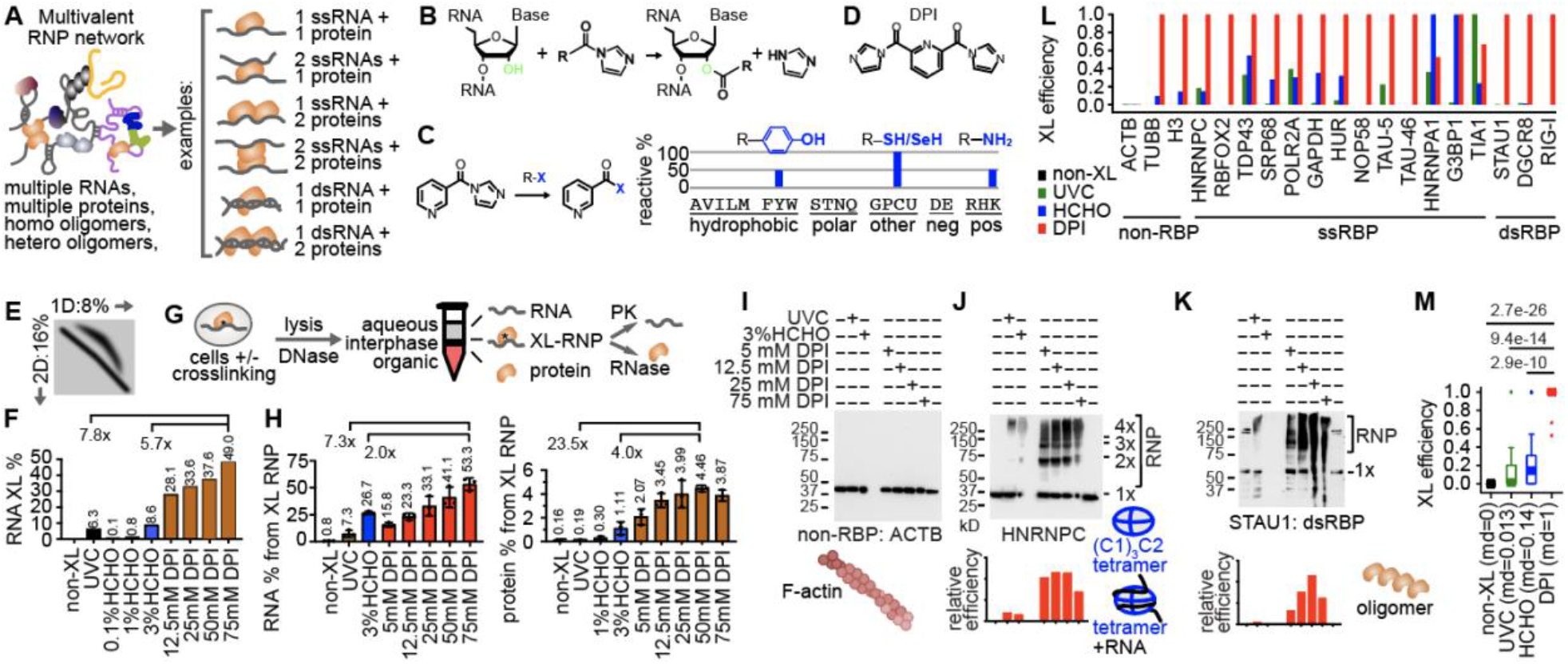
SHARC enables high-efficiency, broad-spectrum capture of multivalent RNP complexes. (A) Schematic of the multivalent RNP network. (B-D) SHARC reaction mechanism. (B) SHARC reagents acylate the RNA 2′-OH group. (C) SHARC reactivity with nucleophilic amino acid side chains, e.g., Tyr, Cys and Lys based on NMR analysis.(D) Chemical structure of dipicolinic acid imidazolide (DPI), the SHARC crosslinker used in this study. (E-F) DD2D gel electrophoresis separates the upper diagonal crosslinked RNAs from non-crosslinked (E), revealing higher RNA-RNA crosslinking efficiency for DPI (F). (G) The aqueous-organic phase separation to quantify RNA-protein crosslinking. Crosslinked RNPs partition to the interphase, separating them from free RNA (aqueous) and free protein (organic).(H) DPI outperforms UVC and HCHO in RNA-protein crosslinking, as measured by the recovered amount of RNA (left) and protein (right). (I-K) Western blot validation of crosslinking efficiency for three proteins. (L) Summary of crosslink efficiencies across 19 proteins. Maximal efficiency of the 4 DPI concentrations was set to 1. (M) Summary of crosslinking efficiencies in box plots. Medians (md) and p-values indicated; two-sided Student’s t-test.

### Efficient and reversible RNP capture by SHARC

To capture multivalent RNP contacts (**Fig. 1A**), we leveraged the RNA 2’-OH crosslinker SHARC, specifically dipicolinic acid imidazolide (DPI) (*15*). In vitro analyses revealed that DPI reacts strongly with nucleophilic residues (Tyr, Cys, Lys) and N-terminal amines, thus enabling the simultaneous capture of RNA-RNA, RNA-protein, and protein-protein contacts (**Fig. 1B-D, Fig. S1A**, supplementary text). Benchmarking against 254nm UVC and formaldehyde (HCHO) revealed that DPI achieves 5.7-to 7.8-fold higher efficiency in RNA-RNA crosslinking and 2.0-to 23.5-fold higher efficiency in RNA-protein capture (**Fig. 1E-H, Fig. S1B**) (*18, 19*). Cumulatively, this represents a 10-to 50-fold improvement over standard methods. Crucially, unlike irreversible UV or inefficiently reversed HCHO crosslinks, SHARC reaction with RNA is fully reversible via mild hydrolysis (**Fig. S1C**).

We benchmarked SHARC across a diverse set of 19 proteins. DPI demonstrated superior or equivalent efficiency for 16 of 18 RNA-binding targets compared to UV and HCHO, while maintaining specificity (**Fig. 1I-M, Fig. S1D**, supplemental text, **table S1**). Notably, only DPI captured the full HNRNPC tetramer with RNA, and crosslinked challenging targets, including RBFOX2, NOP58, and double-strand RBPs STAU1, DGCR8, RIG-I (**Fig. 1J-K**). On average, DPI outperformed HCHO by 7.1-fold and UVC by 76.9-fold, establishing SHARC as a high-efficiency, broad-spectrum chemistry for RNP structure and interaction analysis (**Fig. 1L-M**).

### SHARCLIP captures protein-associated and independent RNA structuromes

To systematically map the dynamic RNP landscape, we designed a split SHARCLIP workflow that disentangles intrinsic RNA folding from protein-associated spatial proximity (**Fig. 2A**). Following crosslinking and antibody-based RNP-enrichment, the RBP-associated path performs on-bead ligation to capture both protein-dependent and independent contacts, while the RBP-independent path ligates directly interacting RNA fragments after proteolysis and DD2D gel separation (**Fig. 2B-C**).

**Fig. 2.**
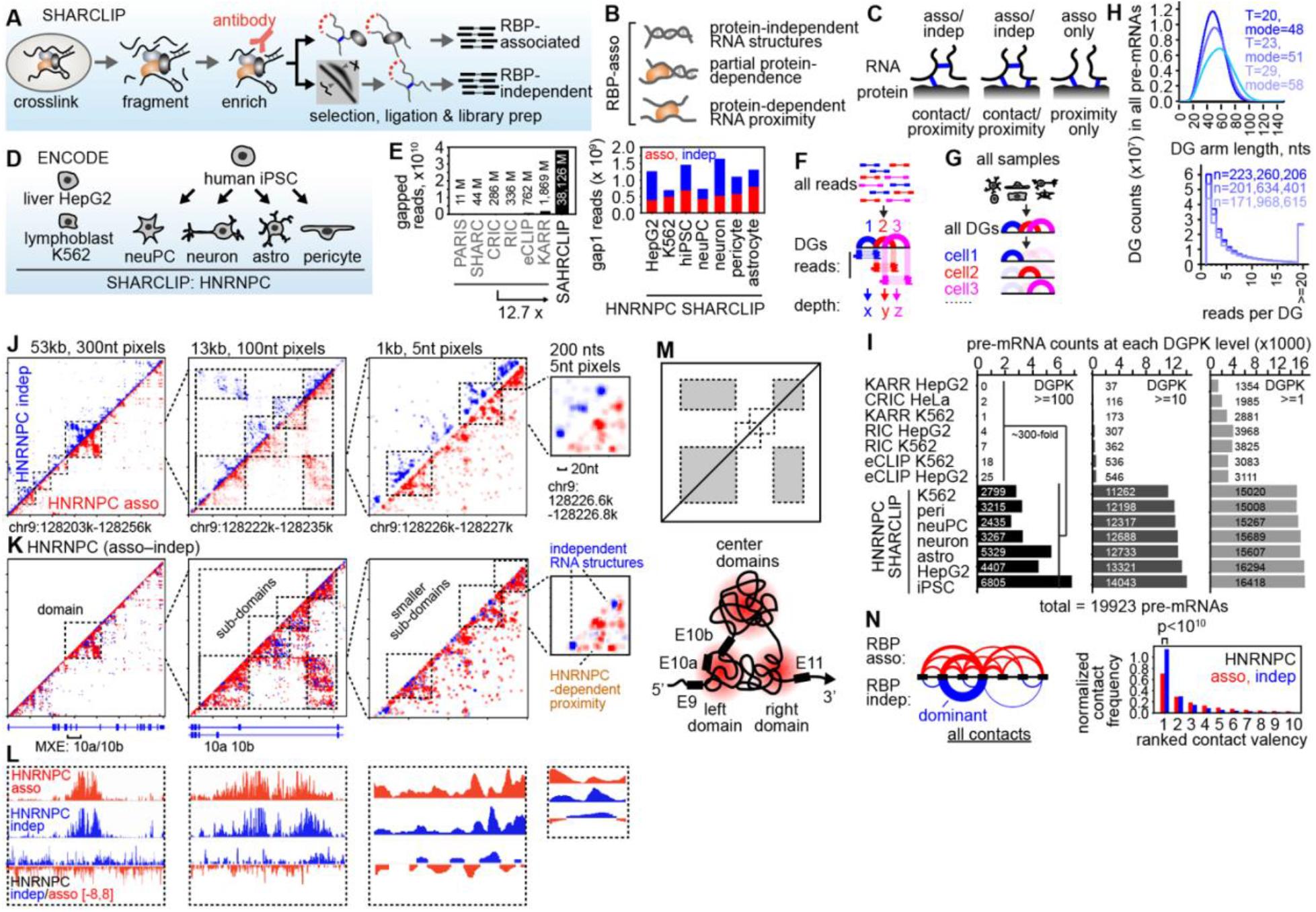
SHARCLIP captures ultra-deep, multi-layered RNA structuromes across 7 cell lines. (A) SHARCLIP workflow. (B-C) The method captures distinct layers of protein-dependent proximity and intrinsic RNA folding. (D-E) HNRNPC SHARCLIP in 7 cell lines generated ~38 billion gapped reads. (F) DG assembly and quantification using CRSSANT-BIRCH. (G) Gapped reads from different samples are assembled into DGs together and then separated for direct and quantitative comparison. (H) Distributions of DG arm length (upper) and read count per DG (lower) for assembled DGs with varying BIRCH threshold (T). (I) pre-mRNA counts at three DGs per kilobase (DGPK) cutoffs. (J-K) Example multi-scale protein-associated and independent RNA conformations captured by HNRNPC SHARCLIP in the DNM1 gene, and their differential. (L) 1D coverage of HNRNPC SHARCLIP data. (M) Overall topology of the 13kb E10a/10b MXE cluster. (N) Global analysis confirms that independent contacts are largely mutually exclusive (low valency), whereas RBP-associated contacts form high-valency regulatory hubs (p < 10^-10^, KS test).

We applied SHARCLIP to HNRNPC, a core component of hnRNPs, across seven diverse cell lineages: HepG2, K562, and an iPSC differentiation series: neural progenitors, neurons, astrocytes, and pericytes (**Fig. 2D, Fig. S2, table S2**). We generated ~38 billion gapped reads, a 12.7-fold increase in depth compared to the combined total of all previous studies (**Fig. 2E, Fig. S3, table S3**) (*18, 20-24*). With 0.7-1.7 billion unique contacts per cell line, we achieved comprehensive coverage of even rare transcripts. Using our CRSSANT-BIRCH pipeline, we assembled these reads into ~200 million unique duplex groups (DGs) (**Fig. 2F-H**) *(25)*, creating a comprehensive structural atlas. Benchmarking confirmed that SHARCLIP significantly outperforms existing methods in efficiency, specificity resolution, and fidelity (**Figs. S4-S5**). The complete structural atlas is accessible via the CRIS database (*26*).

We quantified structural coverage using DGs per kilobase (DGPK), using the mature ribosome (26 helices/kb) and reads sub-sampling as benchmarks for saturation (**Fig. S6A-B**). While previous methods resolved fewer than 25 pre-mRNAs at a high density of DGPK≥100, SHARCLIP captured 2,435-6,805 genes, a ~300-fold improvement (**Fig. 2I**, see **Fig. S6C** for lncRNAs, and **Fig. S6D-E** for examples)). Even at a moderate threshold (DGPK≥10), SHARCLIP covered >11,000 genes, outperforming existing datasets by ~40-fold and approaching transcriptome-wide coverage for the first time.

This depth and resolution enable visualization of RNP topology across vast scales. In the *DNM1* pre-mRNA, we resolved features ranging from ~10kb domains down to 20nt contacts (**Fig. 2J**). The split workflow revealed distinct topological layers: RBP-independent reads defined focal secondary structures, while RBP-associated reads highlighted protein-mediated proximities (**Fig. 2J-K**). These layers allowed us to model the ~13kb exon 10a/10b region as a central domain flanked by interacting side domains (**Fig. 2L**). Interaction partner analysis confirmed that direct contacts exhibit higher mutual exclusivity and lower valency than HNRNPC-associated ones (**Fig. 2M**). Thus, SHARCLIP establishes a new standard for the in vivo structurome, capturing both intrinsic RNA folding and higher-order RNP organization at high resolution.

### Resolving the architecture of the ribonucleosome

Analogous to nucleosomes on DNA (17), the stepwise packaging of nascent RNA into hnRNP particles is critical to processing and export. First visualized in the 1970s as “beads-on-a-web” (27), these particles are assembled on top of the hnRNP A, B, and C tetramers (**Fig. 3A**) (28). The C-tetramer associates with a unitary RNA length of ~235 nts (29, 30). Three C-tetramers with ~700nts RNA assemble into a 19S triad, which recruits A/B tetramers to form 30-40S mono-particles. RNA structures were thought to be interlaced among the protein cores (**Fig. 3B**) (31).

**Fig. 3.**
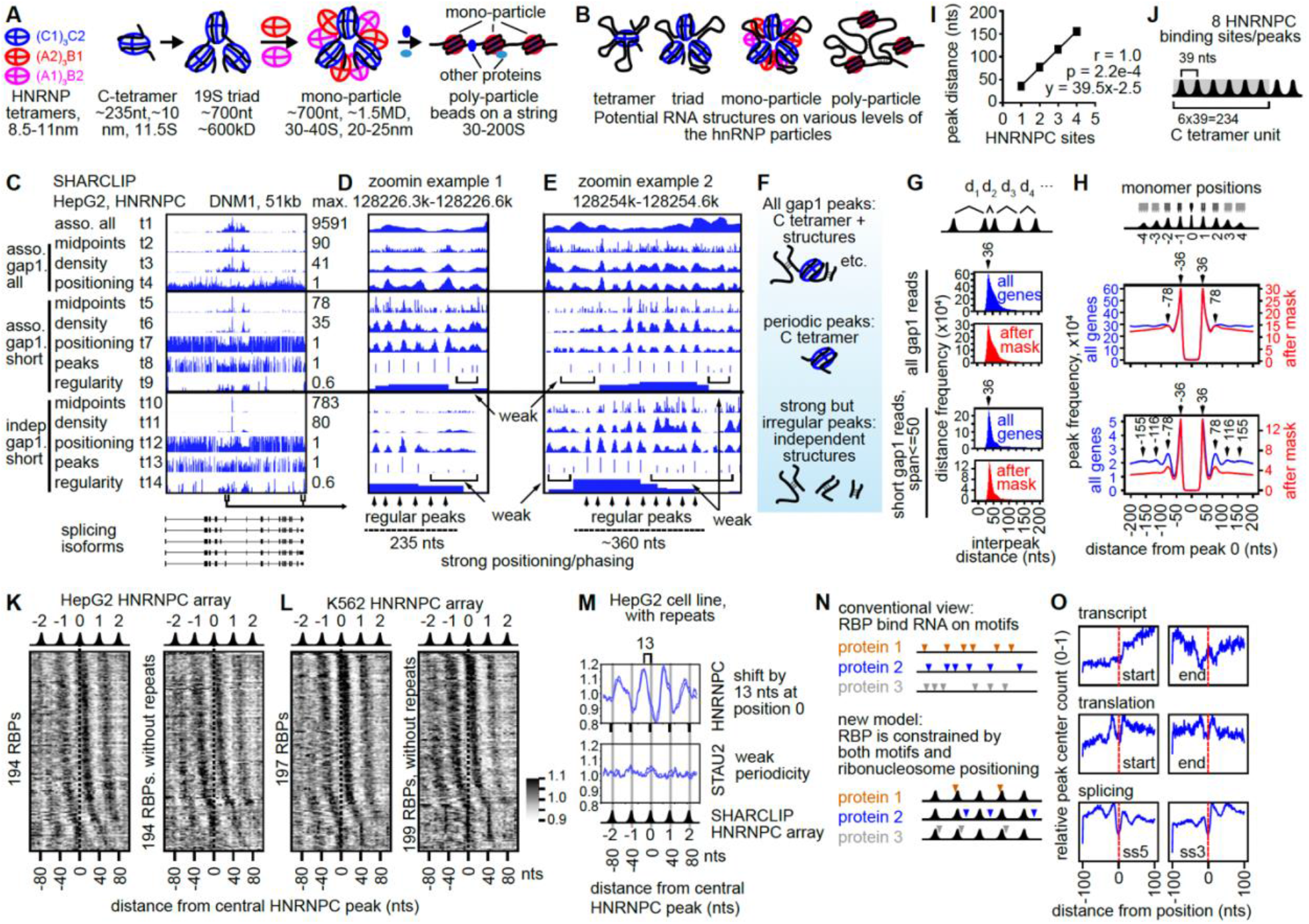
A global map of ribonucleosome positioning, phasing and function. (A-B) Model of hnRNP assembly, showing the RNA-structure-free organization (A) and putatitive locations of RNA structures (B). Estimated sizes are listed in nanometers (nm) and kilo/mega daltons (kD/MD). (C-F) SHARCLIP reveals HNRNPC arrays on the *DNM1* pre-mRNA. (G-H) Global analysis of inter-peak distances reveals a dominant ~39-nt periodicity that persists even after masking repetitive elements. A total of 946845 windows were analyzed which contains 5342304 peaks passing the filter. (I-J) Regression analysis confirms a precise 39-nt repeating unit contributing to the ~235nt unit RNA wrapping each HNRNPC tetramer RNA. (K-M) eCLIP-derived RBP binding sites are phased on SHARCLIP-derived HNRNPC array. (N) A revised model of RBP binding on RNA that considers the core ribonucleosome landscape. (O) HNRNPC SHARCLIP peak average along functional sites, based on all SHARCLIP HNRNPC peaks in HepG2 cells.

However, the cellular organization of these complexes remains controversial. While electron microscopy (EM) and equilibrium binding suggest minimal sequence bias of HNRNPC, UV crosslinking identified a strong uridine motif, leading to a ~165nt spacing model (32). Our reanalysis demonstrates that the 165nt periodicity is an artifact of Alu retrotransposon dominance and absent from the non-repetitive transcriptome (**Fig. S7**, supplementary text). Thus, the true architecture of the ribonucleosome in cells has remained unresolved for over 60 years (33).

We hypothesized that RNA wrapping around C-tetramers generates a distinct topological signature: periodic, protein-dependent proximal contacts (**Fig. S8A**). This contrasts with the non-periodic, variable-distance contacts typical of intrinsic RNA folding. To isolate the periodicity, we adapted algorithms from nucleosome studies (**Fig. S8B-D**) (34). By enriching for short-range contacts within the HNRNPC-associated fraction, we effectively filtered out variable-distance secondary structures to reveal the regular wrapping of RNA around the protein core.

Using the DNM1 locus as a model (**Fig. 3C-F**), we applied a multi-step signal processing pipeline to HNRNPC-associated gapped reads. While raw reads (t1) showed weak periodicity due to the mixture of direct/indirect contacts and heterogeneous boundaries, read midpoints (t2), density (t3) and positioning scores (t4) revealed discrete peaks. Isolating short-range contacts (≤50nt) further improved the signal-to-noise ratio, revealing sharp, clustered peaks arranged in semi-regular arrays (t5-t9). HNRNPC-independent peaks also exhibited periodic subsets despite the weaker signal (t10-t14), suggesting direct tertiary contacts within the RNA wrapped around the C-tetramer.

Peak analysis revealed a precise periodic architecture. Inter-peak distances formed a narrow distribution with a 36-nt mode (**Fig. 3G, Fig. S8E**). Regression analysis corrected the right-skewed distribution, defining a regular spacing of 39 nts (**Fig. 3H-I, Fig. S8F**). This 39-nt phasing provides the missing quantitative link to biochemical data: six periods occupy 234 nts, matching the ~235-nt RNA wrapped on a single C-tetramer (**Fig. 3J**) *(30)*. This architecture mirrors chromatin, where 147 bps of DNA wraps the histone octamer *(34, 35)*. We observed an alternating AU/GC bias with identical periodicity, suggesting that HNRNPC arrays are genetically hardwired, analogous to the ~10.5bp periodicity in nucleosomes (**Fig. S8G-H**) *(34)*. Notably, filtering eCLIP data revealed a HNRNPC arrays but lower periodicity and regularity due to UV crosslink bias and ~50 times lower coverage (**Fig. S8J-L**).

Finally, we assessed whether this landscape regulates other factors. Aligning eCLIP data (22) onto SHARCLIP HNRNPC arrays revealed a surprising differential phasing for diverse RBPs, unexpected based on the known motifs of RBPs (**Fig. 3K-N, Fig. S9**). The HNRNPC UV CLIP sites are also synchronized on the SHARCLIP-derived array, whereas the dsRBP STAU2 showed no periodicity, further confirming the specificity of the array.Furthermore, the arrays exhibit clear phasing relative to splice sites and translation boundaries (**Fig. 3O**) (36). Thus, the C-tetramer in the ribonucleosome imposes a periodic structural constraint on the linear transcriptome. In summary, SHARCLIP reconciles six decades of conflicting evidence to establish the C-tetramer complex as the fundamental, regularly phased unit of cellular hnRNPs.

### Strategies to analyze the structural mechanisms of AS

To determine how HNRNPC-associated and independent structures regulate splicing, we integrated our structural atlas with splicing quantification (**Fig. 4A, data S1, CRIS database)** (*26*). We prioritized functional structures using three key metrics: dominance, the strength of each structure relative to the local total; conservation, specifically focusing on intronic arms; and stability based on model minimum free energy (MFE) (**Fig. 4B-C**) (*37*). We first performed a two-dimensional correlation analysis to link structure to function (**Fig. 4A**). We assessed the global relationship between structural strength and exon inclusion within each cell type, and the correlation between specific conformations and splicing changes across the seven lineages. We further classified structural regulators into three mechanistic categories. Loops are defined as inter-intronic contacts that insulate the exon. Blockers are duplexes where at least one arm directly occludes a core splicing motif. Switches represent dynamic loops and blockers that toggle between mutually exclusive conformations. This framework underlies our mechanistic analysis of alternative splicing (**Figs. 4-6**).

**Fig. 4.**
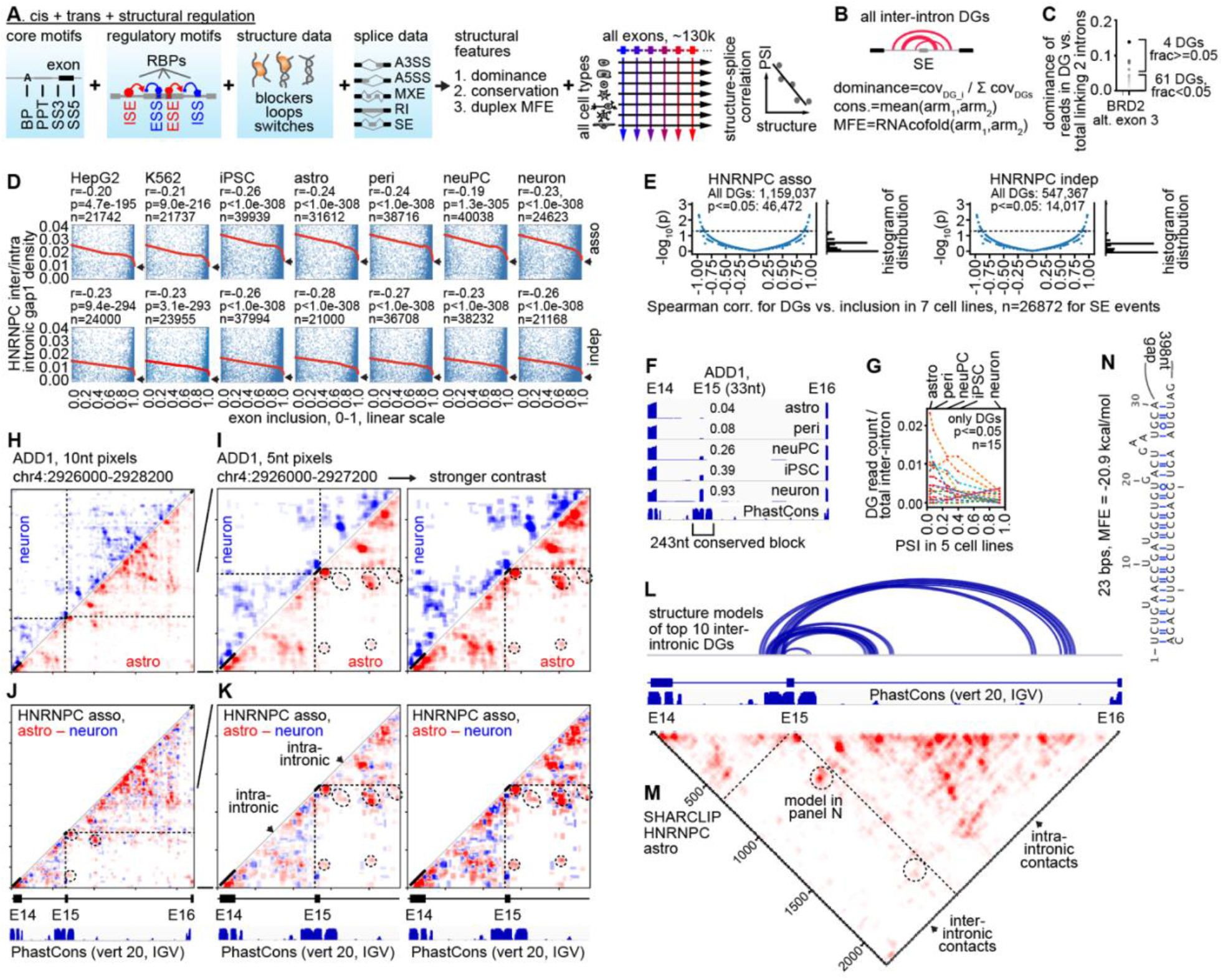
Strategies for the analysis of structural mechanisms and discovery of loops regulating AS. (A) Strategy to identify structural regulators of splicing. ISE, ESE, ISS, ESS: intronic or exonic splicing enhancers or silencers.PSI: percent spliced in. (B) Calculation of structure features using inter-intron loops and a skipped exon (SE) as an example. (C) Example distribution of all fraction values for all loop DGs around the BRD2 alternative exon 3. (D) Spearman correlation between inter-intron gapped reads and exon inclusion levels. Only junctions with >=10 inclusion + skipping reads are used. Red line: LOWESS smoothing. (E) Spearman correlation of normalized counts of each inter-intronic DG vs. the inclusion levels of the associated exon. (F) Differential skipping of ADD1 E15. (G) Correlations between 15 DGs and inclusion levels across the 5 cell types. (H-K) Individual and differential heatmaps of HNRNPC-associated gapped reads between astrocytes and neurons around E15. (L-M) Structure models of top 10 inter-intronic DGs (L) overlapped on the SHARCLIP heatmap (M). (N) Structure model of the strongest loop.

### Inter-intronic loops as splicing insulators

Inter-intronic interactions have the potential to sequester exons by acting as structural insulators. To map these features, we analyzed 27,149 skipped exon (SE) events with variable inclusion across seven cell lines. From ~1.5 million candidates, we filtered for structural dominance (≥5%) and deep conservation (phastCons≥0.9), yielding ~77,000 and ~13,000 high-confidence loops, respectively (**Fig. S10A-B, data S1, CRIS database**) (*26*). Correlation analysis revealed a global negative association between loop strength and exon inclusion (**Fig. 4D**). This relationship follows a non-linear threshold model, where the emergence of loops acts as a sharp brake on exon inclusion (**Fig**.

**4D**, arrows, and **Fig. S10C**). At the event level, we identified 46,472 DGs (4.0% of total) where structural intensity significantly anticorrelates with inclusion (**Fig. 4E**).

As an example, we highlight *ADD1*, a cytoskeleton gene linked to intellectual disability (38). Exon 15 (E15) is flanked by highly conserved regions and displays lineage-specific skipping (**Fig. 4F**). We identified 15 DGs associated with E15 regulation, 12 of which exhibited the expected negative correlation (**Fig. 4G**). Structural mapping revealed that in astrocytes, where E15 is skipped, the exon is sequestered by a dense network of inter-intronic loops, whereas the region remains structurally accessible in neurons, where E15 is included (**Fig. 4H-K**). Modeling confirmed these loops form stable duplexes (**Fig. 4L-N, Fig. S10D**), establishing inter-intronic looping as a pervasive mechanism of negative splicing regulation.

### Splicing modulation via steric blocking

Stable RNA structures can sterically occlude core splicing motifs to inhibit exon recognition (**Fig. 5A-B**). We identified ~3.7 million DGs in ~14,900 genes, corresponding to ~1.7 million unique blockers with high folding stability (**Fig. 5C, Fig. S11A-F, data S2-S5, CRIS database**) (26). We further defined a high-confidence subset of ~148,000 blockers where the non-blocking intronic arm is evolutionarily conserved (phastCons≥0.9). Globally, blocker strength exhibits a robust negative correlation with exon inclusion for SE and MXE (**Fig. 5D**). In contrast, A5SS, A3SS and RI lacked simple correlations, likely due to the tight spatial constraints of these events (**Fig. S11G**). While insufficient variation in inclusion levels can mask these trends, we identified dozens to thousands of events where structural occlusion significantly predicts splicing outcomes (**Fig. 5E**).

**Fig. 5.**
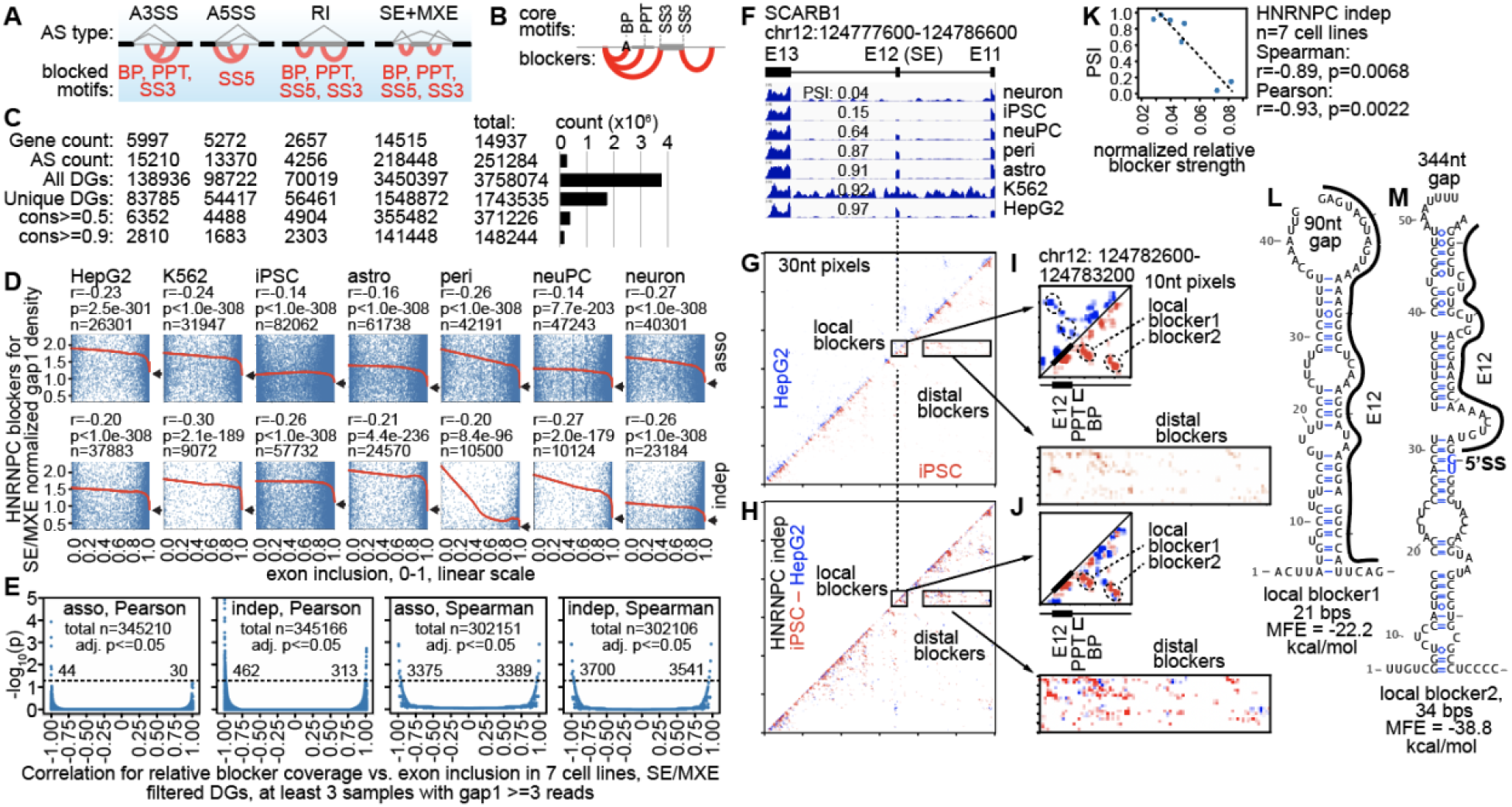
SHARCLIP identifies structural blockers of alternative splicing (AS). (A) Possible blockers for all AS types. (B) Zoom-in view of blockers. (C) Global discovery of structural blockers. Cons: vertebrate 100-way phastCons score. (D) Correlation between inclusion levels and blocker strengths. Blocker DG coverage was normalized against the average of two neighbor exons for all SE events with average exonic coverage ≥1. Spearman’s rank correlation coefficients, p-values, and numbers of SE events are listed. Red curve: LOWESS smoothing. Arrows point to the right ends of the LOWESS curves. (E) Pearson and Spearman correlations of HNRNPC SHARCLIP asso or indep blocker gapped reads vs. inclusion levels for SE. X-axes: correlation coefficients. Y-axes: −log10(p) after Benjamini-Hochberg correction. Numbers of DGs with significantly negative and positive correlations are listed above the dotted lines (adj. p<=0.05). (F) A differentially skipped exon in SCARB1 across 7 cell lines. (G-H) HNRNPC SHARCLIP indep. gapped reads are plotted as separate or differential heatmaps. (I-J) Two major groups of local and distal blockers. (K) The correlation between PSI values for the E12 and blocker strength normalized against E11+E13 coverage are tested using Spearman and Pearson. (L-M) Secondary structure models for the two local blockers.

To demonstrate this mechanism, we analyzed *SCARB1*, a lipoprotein receptor gene linked to coronary artery disease (39). SCARB1 exon 12 (E12) shows strong differential inclusion, ranging from 4% to 97% (**Fig. 5F**). SHARCLIP revealed a dense set of blockers specifically in iPSCs (where E12 is excluded). These include both distal (~4kb range) and high-intensity local structures (**Fig. 5G-J**). The strength of these blockers varies by ~3 times among the cell lines, showing a strong anti-correlation with exon inclusion (**Fig. 5K**). Structural modeling indicates that the dominant local blockers form two stable helices to sequester E12 (**Fig. 5L-M**). This suggests *SCARB1* splicing is enforced by a cell-type-specific conformation that sterically occludes the exon to drive skipping.

### Structural mechanisms determine MXE choice

The human genome contains ~230 high-confidence MXE clusters that function as biological “exclusive OR” logic gates (**Fig. 6A**). While critical for development, cytoskeletal organization and synaptic transmission, the regulation of these clusters remains poorly understood (**Fig. S12A**) (40). Canonical models, e.g., NMD, incompatible splice sites, and steric hindrance explain only a few events (**Fig. 6B**) (41). Beyond isolated examples like *Dscam* and *ATE1*, it was unknown if direct structural switching is a general paradigm (42-44).

**Fig. 6.**
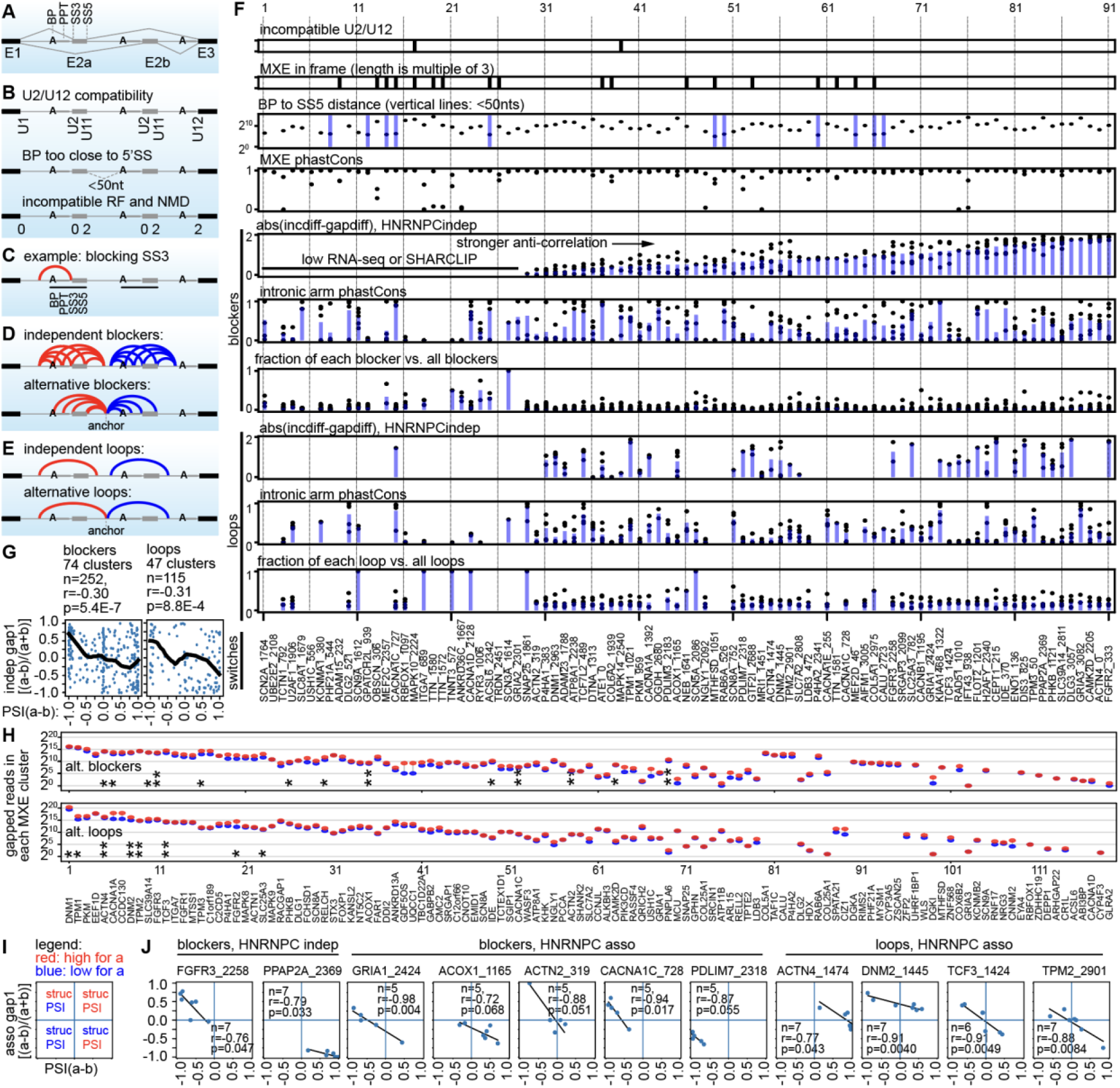
Discovery of structural mechanisms regulating MXEs. (A) Organization of a typical MXE cluster and core splicing motifs. (B) Three linear mechanisms that enforce mutual exclusivity. (C) Example blocker for the SS3. (D) Independent blockers for BP, PPT, SS3 or SS5, with anchor arms, or dock sites, in the three neighbor introns (upper panel). Alternative blockers that share the anchor arms are potential structural switches (lower panel). (E) Independent loops covering two MXEs (upper panel). Alternative loops with shared anchor arms are potential structural switches (lower panel). (F) The mechanism matrix for 91 MXE clusters with RNA-seq and SHARCLIP data. The abs(incdiff-gapdiff) tracks were computed the same way as in panel G, and high values indicate strong anti-correlations. Incdiff is PSI difference between a and b, or PSI(a-b). Gapdiff is normalized gapped read coverage difference between a and b, or (a-b)/(a+b). Each dot represents one cell lines. For blocker and loop intronic arm phastCons, top 5 DGs are shown. (G) Pearson correlation between the differential PSI values (MXE isoform a vs. b), and relative differential gapped read counts normalized to the range of [-1,1]. Each dot is one MXE cluster in one cell line. Thick black lines: LOWESS smoothing. (H) Dumbbell plot of structural switches for each MXE cluster. Red and blue dots represent the blocking gapped read counts from all 7 cell lines for the two exons in each cluster. Asterisks indicate MXEs with biased quadrant distributions (*) or negative correlations (**) between gapped read count and exon inclusion levels. (I) Diagram explaining the 4 quadrants. (J) Example negative correlations between blockers/loops and exon inclusion levels. Number of cell lines with quantifiable structures and splicing, Pearson correlation coefficients and p values are listed for each MXE cluster.

We hypothesized that alternative blockers and loops physically enforce this exclusivity (**Fig. 6C-E**). Leveraging our quantitative structural atlas across 91 conserved MXE clusters, we identified specific blockers and loops for 64 clusters (70%) (**Fig. 6F, Fig. S12B-C**). These structural regulators are often dominant, deeply conserved, and exhibit a global anti-correlation with exon inclusion (**Fig. 6F-G, Fig. S13D-E**). Expanding this analysis to all structural data, we discovered switches in 119 clusters, recovering *ATE1* as a validation of our approach (**Fig. 6H, Fig. S13A-C, data S6-S7, CRIS database**) (*26*). At the individual level, quadrant analysis of MXEs with low inclusion variability identified 23 clusters regulated by alternative blockers or loops (**Fig. S13F-H**). Furthermore, among highly variable MXEs, we identified 11 clusters where inclusion anticorrelates dynamically with structural changes across the cell lines (**Fig. 6I-J**). Together, these data demonstrate that structural switches are not rare exceptions but a prevalent mechanism that enforce mutually exclusivity.

### A thermodynamic switch of *TCF3* MXEs in development

To validate the structural switch models, we focused on *TCF3*, a master stem cell regulator linked to leukemia and immunodeficiencies *(45-48)*. The E18a and E18b MXEs produce functionally distinct isoforms via a developmental switch that has remained mechanistically obscure for 40 years *(49, 50)*. SHARCLIP revealed a dynamic landscape where a conserved intronic control region (ICR) acts as a structural anchor. In iPSCs, the ICR preferentially forms duplexes with E18b, blocking it to favor E18a, whereas in pericytes, the conformation flips to block E18a (**Fig. 7B-E, Fig. S14A-D**). This “exclusive OR” logic was confirmed by a global negative correlation between blocker strength and inclusion across five cell types (**Fig. 7F-J**).

**Fig. 7.**
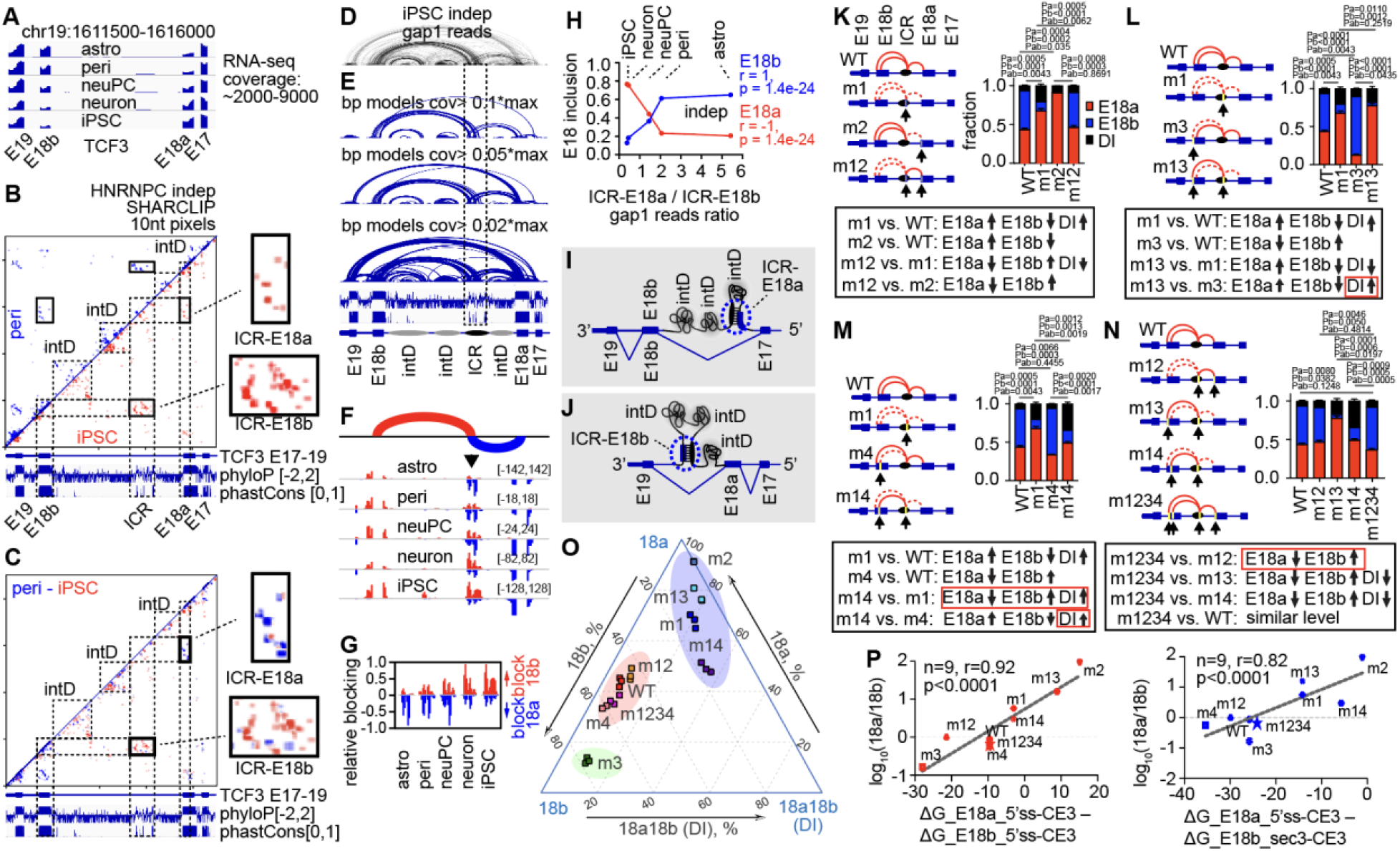
Developmentally dynamic structural switches control the TCF3 MXEs. (A) RNA-seq quantification of exon inclusion. (B-C) SHARCLIP gapped reads were plotted in separate (B) and differential (C) heatmaps. intD: internal domains. ICR: intronic control region. (D) SHARCLIP gapped reads were plotted as arcs. (E) Base-pair models for DGs with relative coverage above each cutoff: 0.1, 0.05, or 0.02 times the max DG in this region. (F) Intronic blockers against E18a and E18b. (G) Zoom-in view of the mutually exclusive ICR blockers against 18a and 18b. (H) Scatter plots of E18 inclusion levels vs. the ratios of gapped reads of the ICR blockers. Coefficients (r) and p values: Spearman rank correlation. (I-J) Model of the ICR-E18a-E18b switch. (K-N) Quantification of exon usage in wildtype (WT) and mutant minigenes. DI: double inclusion. Solid red curves: stable blockers, WT or after compensatory mutations. Dashed red curves: mutation-disrupted blockers. Two-sided t-test p values were calculated for 18a (Pa), 18b (Pb) and DI (Pab). Red boxes: minor deviations. (O) Ternary plot of inclusion levels in the 9 minigenes. (P) Scatter plots of the log frequency of 18a/18b differential inclusion levels vs. differential blocker MFEs, linear correlation coefficients and p values.

We proved necessity and sufficiency using a minigene reporter (**Fig. S14**). Disrupting the ICR anchor (m1) weakened mutual exclusivity, increasing double inclusion. The increase of E18a and decrease of E18b is consistent with the transcription direction and proximity of E18a to ICR (**Fig. 7K**). We further dissected the switch by mutating specific arms. Disrupting the E18a-blocker (m2) increased E18a inclusion, while disrupting the E18b-blocker (m3, m4) shifted splicing toward E18b (**Fig. 7K-N**). Most importantly, compensatory mutations that restored base-pairing despite altered sequence rescued wild-type splicing patterns (m12, m13, m14, and m1234, **Fig. 7K-O, Fig. S15**, supplementary text). Finally, we found that the thermodynamic stability (MFE) of these mutants linearly predicts the log-ratio of exon inclusion (**Fig. 7P, Fig. S14H**). This relationship aligns with the Boltzmann distribution, p ~ exp(-E/(k_B_T)), providing a quantitative physical basis for this biological decision. Therefore, the TCF3 MXEs are physically enforced by a thermodynamic switch.

### Structure-guided ASO design for therapeutic splice switching

Pathogenic mutations are linked to splicing by many mechanisms, such as weakening splice sites, creating cryptic exons, or mutating amino acids in alternative exons. Our discovery of structural regulators offers new therapeutic avenues for these defects. Mutually exclusive exons (MXEs) are particularly enriched in pathogenic mutations, presenting a unique opportunity to rescue phenotype by forcing a switch to the homologous healthy isoform (*40, 51, 52*). We developed a generalized rational framework to design ASOs that modulate splicing by targeting RNA structures rather than linear motifs. Our approach employs three distinct modalities: duplex breaker ASOs to competitively open repressive structures, and triplex and three-way junction (3WJ) forming ASOs to stabilize weak structures (**Fig. 8A-C**) (*53*). To demonstrate broad applicability, we surveyed human genetic data and identified 35 genes where MXE mutations impact more than 10 organs and systems, primarily brain, heart, and skeletal muscle (**Fig. 8D-G, table S4**) (*54*). These MXEs control ion channels, synaptic transmission, structural proteins, and metabolism, and the mutations cause epilepsies, neuromuscular disorders, and other diseases. We selected 12 high-value targets, prioritizing structures based on dominance, stability, and splicing correlation (**Fig. 8E-F**).

**Fig. 8.**
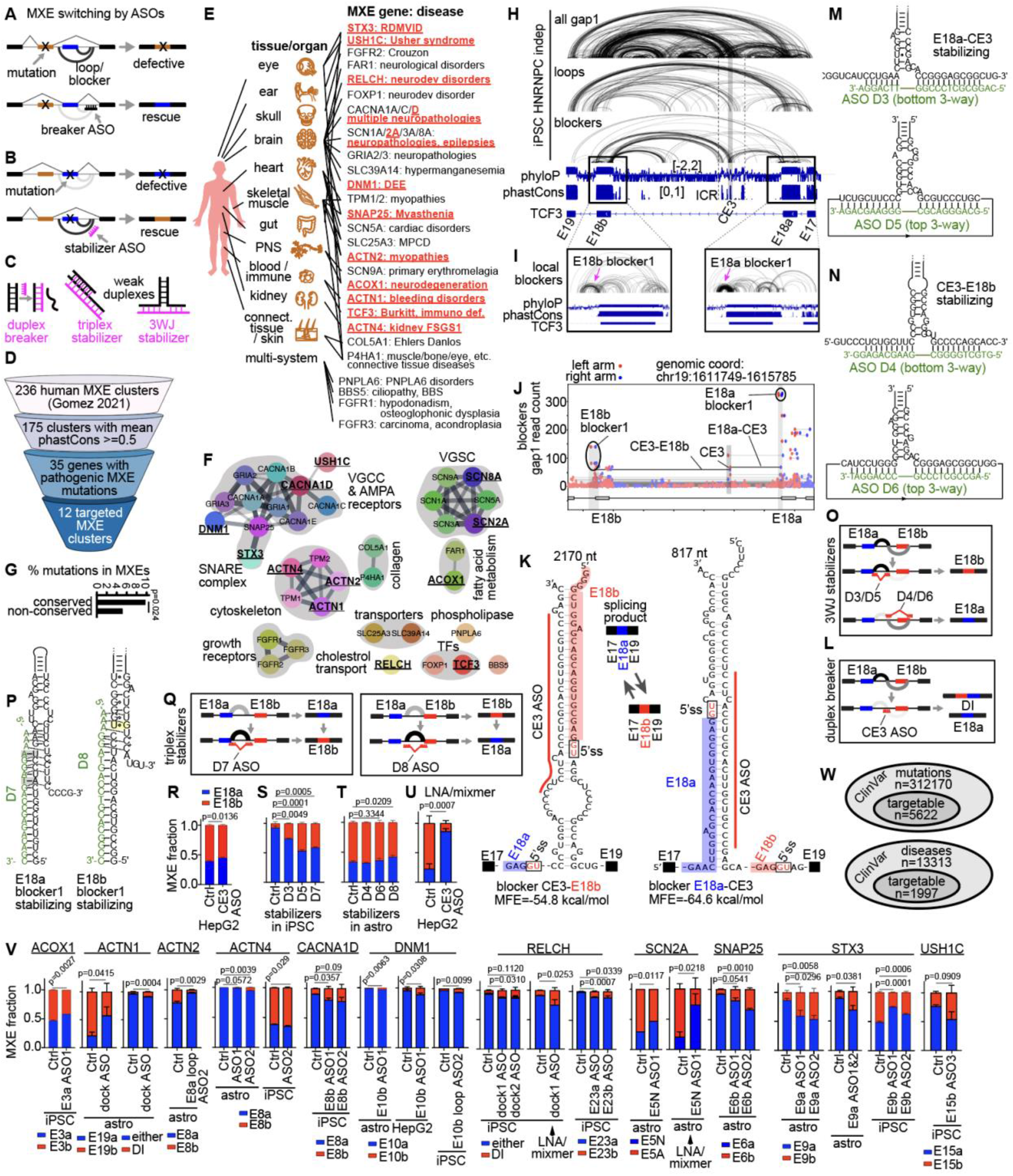
Rational design of structure-targeting ASOs for splice switching. (A-B) Schematic of ASOs designed to either break or stabilize specific conformations to induce isoform switching. (C) Three structural targeting modalities.(D) Identification of targetable MXEs. (E-F) Genetic disorders with pathogenic mutations mapped to MXEs and potentially amenable to structure targeting. Genes to be targeted are highlighted. RDMVID: retinal dystrophy and microvillus inclusion disease. DEE: developmental and epileptic encephalopathies. MPCD: mitochondrial phosphate carrier deficiency. VGCC/VGSC: voltage-gated calcium/sodium channels. (G) 97 and 13 single amino acid mutations map to 1003 conserved and 256 non-conserved residues in the MXEs. Chi-square test, p=0.024. (H-I) Structural loops and blockers around the TCF3 MXEs. (J) Quantification of blockers, highlighting E18a blocker 1, E18b blocker 1, CE3-E18b and E18a-CE3. (K-L) Structure models of the switch and the location of the CE3 ASO. DI: double inclusion. (M-O) Design of the 3WJ stabilizing D3-D6 ASOs targeting the E18a-CE3 and CE3-E18b blockers. (P-Q) Design of the triplex stabilizer D7 and D8 ASOs targeting the two local blockers. (R-T) The MOE ASOs were transfected into cells and MXE fractions were measured by PCR and restriction enzyme digestion. p values: two-sided Student’s t-tests. (U) Effect of the locked nucleic acid (LNA)/mixmer CE3 ASO on the TCF3 MXEs. (V) Summary of structure-targeting ASOs for 11 additional MXE-containing genes. All ASOs are MOE modified except the three LNA/mixmers for TCF3 CE3, RELCH dock1 and SCN2A E5N ASO1. (W) Interaction of SE blockers, ClinVar pathogenic and likely pathogenic mutations, and spliceAI predicted hypomorphic variants with acceptor/donor loss in the range of 0.2-0.8. Total mutations and diseases, as well as targetable subsets using structure manipulation are listed in Venn diagrams, which are not plotted to scale.

We first focused on TCF3, where mutations cluster in E18b (**Fig. S16**). Comprehensive structure analysis prioritized four structural regulators (**Fig. 8H-J**). We first targeted the ICR switch and local blockers using 2’-methoxy-ethyl (MOE)-modified ASOs (**Fig. 8K-Q**). As expected, breaking the switch anchor favored E18a inclusion, and triplex and 3WJ ASOs successfully stabilized specific conformations to force the opposite splice choice (**Fig. 8R-T**).Notably, optimizing chemistry to locked nucleic acid (LNA)/mixmers drastically enhanced potency compared to MOE (**Fig. 8U**). Expanding beyond *TCF3*, we validated this platform across 11 additional disease-relevant genes. In every gene, structure-targeting ASOs induced significant isoform switching (**Fig. 8V, Figs. S17-27**, supplementary text). LNA/mixmer designs consistently outperformed MOE, particularly for targets like *RELCH* and *SCN2A*. These results validate the newly discovered blocker/loop/switch mechanisms and establish SHARCLIP as a foundation for rapid and rational design of structure-targeting RNA therapeutics. In addition to MXEs, this strategy extends to broader splicing defects, such as rescuing hypomorphic mutations by disrupting splicing inhibitors, as already demonstrated in the SMN2 case (*55*). To quantify this scope, we analyzed ClinVar pathogenic and likely pathogenic variants using SpliceAI predictions (*54, 56*). We identified 5,622 splice-weakening variants (1.8%) spanning 1,997 diseases (15.0%) that are also inhibited by blockers, and thus potentially targetable by breaking the blockers to at least partially rescue the weak splicing (**Fig. 8W**). These numbers are conservative estimates given we restricted our analysis to core motif blockers.

## Discussion

The development of SHARC reagents represents a fundamental expansion of the RNA chemical biology toolkit (*15*). Unlike traditional crosslinkers (UV, HCHO) that suffer from low efficiency, bias, and irreversible damage, SHARC chemistry achieves high-efficiency, reversible capture of RNA-RNA, RNA-protein, and protein-protein contacts (reversibility only applies to RNA). Taking advantage of this versatility, we establish SHARCLIP as a new standard for in vivo structural analysis, delivering superior resolution and the critical ability to distinguish direct physical contacts from broad spatial proximity (*12*).

These innovations culminated in the first ultra-deep atlas of human nuclear RNA structuromes. We built the first molecular map of cellular ribonucleosomes, revealing a 39-nt periodic architecture that packages nascent RNA, reminiscent of the well-studied nucleosomes. Future mapping of A/B group proteins and other RBPs will be essential to fully understand the interplay between the hardwired and plastic structural landscapes.

By profiling diverse cell lineages, we demonstrate that RNA folding is cell type-dependent, providing a foundational resource for splicing biology. We identified structural regulators for >70% of protein-coding genes, classifying them into a mechanistic framework of blockers, loops, and switches. We resolved the mechanisms that control the TCF3 MXEs, proving that mutual exclusivity is enforced by a genetically hardwired, thermodynamical switch. Future expansion to diverse human tissues and developmental stages will be critical to fully map the complete structural basis of RNA processing in physiology and disease. While we focused here on splicing, the structural code likely governs other processing and functional steps of the RNA life cycle, such as editing, polyadenylation, localization and translation. Although our current classification is foundational, additional mechanisms, such as bridging, and regulatory motif blocking, can be further explored (*57*). The immense volume of data necessitates the development of deep learning models to predict regulation from sequence and structure, moving the splicing code from 1D to 3D.

Finally, this work opens a new frontier in precision medicine: structure-guided RNA therapeutics. We provide proof-of-concept that splicing-related pathogenic mutations can be rescued by rationally targeting structural regulators with ASOs. Even though we focused on MXEs, other AS events, which together cover >90% of human protein-coding genes can be targeted in similar manners. This approach is faster and more precise than traditional oligo walking, offering a rapid-response platform for rare genetic disorders that together impact ~10% of the human population. This structure-first approach is not limited to ASOs; the stable pockets and helices identified here represent prime targets for small molecules, another challenging but promising frontier. Together, SHARCLIP bridges the gap between static genetics and dynamic structural biology, unlocking the transcriptome as a vast library of druggable targets.

## Acknowledgments

We thank members of the Lu lab, including Wilson Lee, Aishi Liang, and Fengjia Chen for discussion and assistance, members of the Guo lab, including In Kyu Yang, Justin Pi, Brian Nguyen, Alina Tong, and Scott Hale for ASO synthesis. We acknowledge support from the USC Center for Advanced Research Computing.

## Author contributions

Conceptualization: Z.L.; Methodology: J.B., K.L., W.Z., G.S., W.A.V., J.C., F.G., Z.L.; Investigation: J.B., K.L., W.Z., G.S., W.A.V., J.C., F.G., and Z.L.; Visualization: J.B., K.L., M.Z., G.S., Z.L.; Funding acquisition: Z.L.; Project administration: Z.L.; Supervision: J.C., F.G., Z.L.; Writing - original draft: J.B., K.L. and Z.L.; Writing - review and editing: all authors.

## Data, code and materials availability

Raw sequencing data were submitted to GEO under accession numbers GSE317330 (SHARCLIP) and GSE316843 (bulk RNA-seq). Public sequencing datasets used in this study were acquired from GEO under the following accession numbers: GSE127188 (RIC-seq), GSE210582 (CRIC-seq), and GSE166155 (KARR-seq). RBP eCLIP data were accessed from the ENCODE Project (https://www.encodeproject.org/). Processed structure data are shared in the CRIS database, https://whl-usc.github.io/cris/home. Code and scripts for data analysis are available in Github: https://github.com/zhipenglu/CRSSANT and https://github.com/zhipenglu/SHARCLIP. Additional materials and information required to reanalyze the data reported in this paper is available upon request.

## List of Supplementary Materials

Materials and Methods

Supplementary text

Figs. S1 to S27

Tables S1 to S6

Data S1 to S7

References

